# Differential proportionality - a normalization-free approach to differential gene expression

**DOI:** 10.1101/134536

**Authors:** Ionas Erb, Thomas Quinn, David Lovell, Cedric Notredame

## Abstract

Gene expression data, such as those generated by next generation sequencing technologies (RNA-seq), are of an inherently relative nature: the total number of sequenced reads has no biological meaning. This issue is most often addressed with various normalization techniques which all face the same problem: once information about the total mRNA content of the origin cells is lost, it cannot be recovered by mere technical means. Additional knowledge, in the form of an unchanged reference, is necessary; however, this reference can usually only be estimated. Here we propose a novel method where sample normalization is unnecessary, but important insights can be obtained nevertheless. Instead of trying to recover absolute abundances, our method is entirely based on ratios, so normalization factors cancel by default. Although the differential expression of individual genes cannot be recovered this way, the ratios themselves can be differentially expressed (even when their constituents are not). Yet, most current analyses are blind to these cases, while our approach reveals them directly. Specifically, we show how the differential expression of gene ratios can be formalized by decomposing log-ratio variance (LRV) and deriving intuitive statistics from it. Although small LRVs have been used to detect proportional genes in gene expression data before, we focus here on the change in proportionality factors between groups of samples (e.g. tissue-specific proportionality). For this, we propose a statistic that is equivalent to the squared *t*-statistic of one-way ANOVA, but for gene ratios. In doing so, we show how precision weights can be incorporated to account for the peculiarities of count data, and, moreover, how a moderated statistic can be derived in the same way as the one following from a hierarchical model for individual genes. We also discuss approaches to deal with zero counts, deriving an expression of our statistic that is able to incorporate them. In providing a detailed analysis of the connections between the differential expression of genes and the differential proportionality of pairs, we facilitate a clear interpretation of new concepts. The proposed framework is applied to a data set from GTEx consisting of 98 samples from the cerebellum and cortex, with selected examples shown. A computationally efficient implementation of the approach in R has been released as an addendum to the propr package.^1^

## 1 Introduction

Normalization techniques for transcriptome sequencing data continue to be of high interest to the data analysis community (e.g. see (Dillies *et al.*, 2013) for a review and (Lun *et al.*, 2016) for a recent example in single-cell RNA-seq). For sample normalization between entirely different conditions, however, ever more sophisticated techniques cannot close the knowledge gap that is of a principal nature: the total mRNA content of the cells of origin is unknown and can only be obtained with an appropriate ‘absolute’ technique.

It has been argued that normalizations can be avoided by performing a log-ratio transformation of the data (Fernandes *et al.*, 2013; Lovell *et al.*, 2015). Such data transformations, however, depend on the reference that is used. The danger here is that the resulting transformed data is ultimately interpreted in a gene-wise fashion. Interpreting log-ratio transformed expression data as referring to gene abundances (instead of ratios with respect to a given reference) runs into the exact same problems as using normalizations (Erb & Notredame, 2016). It effectively means that the log-ratio transformation is seen as a normalization (that has, as it were, an additional aura of technical sophistication). The only way out of this dilemma seems to be to let go of the gene-wise perspective entirely and instead consider ratios as the basic objects of interest. Although some information will remain hidden this way (such as the true differential gene expression between absolute abundances), the remaining signal will be inherently unbiased.

Here we propose a formal framework for understanding *differential ratio expression*, a change in the ratio of abundances between experimental groups. In doing so, we show that techniques developed for the analysis of the differential expression of genes (e.g. methods known from the limma/voom approach (Smyth, 2004; Smyth, 2005; Law *et al.*, 2014) apply to the analysis of differential ratios as well. This seems intuitive when considering gene ratios as depicted in Figure 1D: an identical picture could be obtained using read counts of a differentially expressed gene instead of gene ratios as shown. However, the interpretation of differential ratios differs considerably.

**Figure 1:**
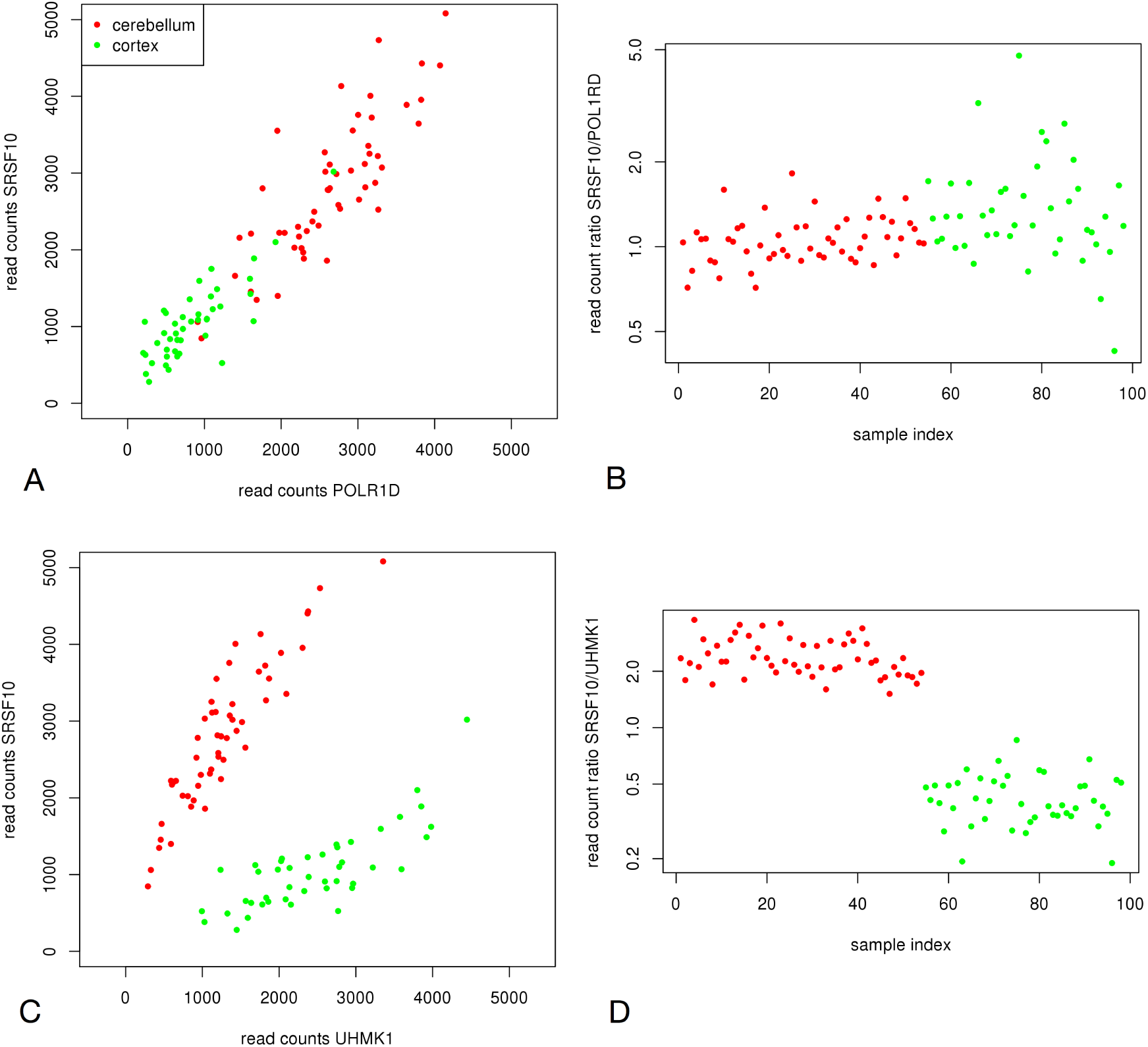
Constant and changing ratios across 98 samples from two tissues: (A) Scatter plot of two genes having an approximately constant read count ratio across all samples (i.e. proportional genes). (B) Ratio plot of the same two genes as in panel A. Although panel A suggests their differential expression, ratios are unable to reveal it. (C) Example of *differentially proportional* genes. Their correlation appears to be about equally strong in both tissues, but the slope of their linear relationship changes between the tissues. (D) Ratio plot of the same two genes as in panel C. The tissue-specific proportionality factors can be detected clearly, and the picture suggests that conventional methods of differential gene expression can be applied to ratios as well.

First, we must consider what it means for a gene ratio to remain unchanged across all sample data. The answer is that the two genes change in the same way (or otherwise remain both unchanged). Figure 1A shows this case in a scatter plot of the read counts for two genes (a splicing factor and a polymerase subunit). Note that although the gene ratio may remain the same, the genes themselves could have joint differential expression. Such gene-wise differential expression is not detected by the ratio approach: although the two genes appear differentially expressed between the tissues, their approximately constant ratio, as shown in Figure 1B, does not reveal this. However, without knowing absolute mRNA abundances, genes may appear differentially expressed only as an artifact of their relative nature.

Second, we must consider what it means for a gene ratio to differ between experimental groups. Figures 1C and 1D shows an example of tissue-specific gene ratios. Here, the two genes (the same splicing factor as before and a kinase) are correlated in both tissues (with a similar strength of correlation), but with different slopes. This means their proportionality factor is tissue-specific (i.e. they have *differential proportionality*). In terms of biochemistry, this could indicate a change in the stoichiometry of the protein products resulting from these mRNAs. Preliminary GO-category enrichment analyses support this view, showing that differentially proportional pairs often contain genes that form protein complexes like those involved in transcription or ribosomal activity.

Current standard methods are not tailored to infer differentially proportional pairs (c.f., Figure 3), although a special class of them, involving receptor subunits in the human brain, has been found by considering time-dependent correlations (Bar-Shira *et al.*, 2015). One method, differential correlation (Tesson *et al.*, 2010), is concerned with differential correlation coefficients, but not with the differential slopes of linear relationships. Importantly, current methods always include a normalization step that–in the best case scenario–introduces extra noise, thus reducing efficacy compared with a method that picks up such signals directly.

**Figure 2:**
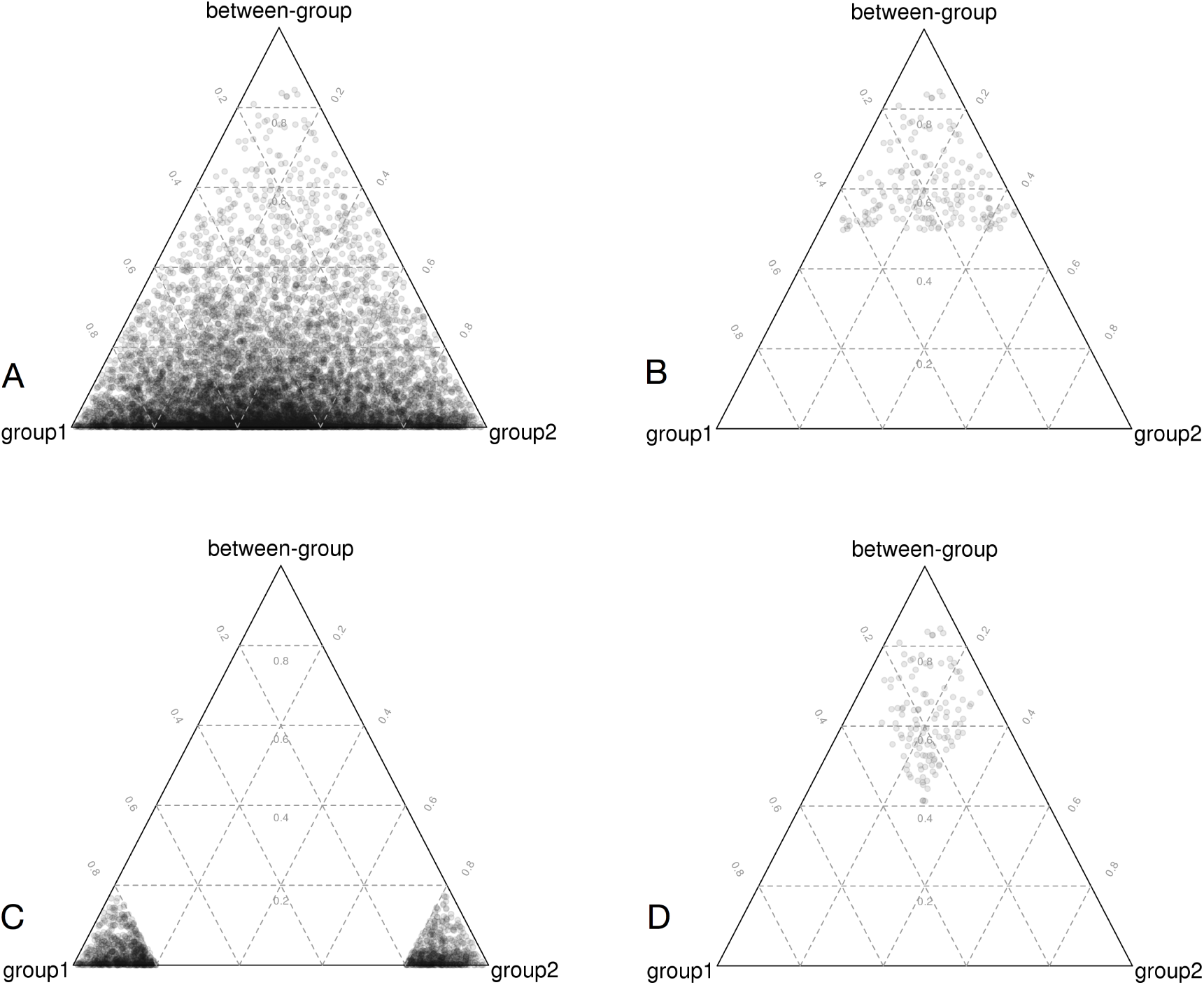
Decomposition of log-ratio variance into (weighted) group variances and between-group variance shown in ternary diagrams. Data from our example from GTEx (group 1 cerebellum, group 2 cortex) are shown. For better visibility, a subset of 10,000 randomly sampled gene pairs were selected. (A): The 10,000 dots corresponding to LRVs of each gene pair. (B): Gene pairs fulfilling *ϑ <* 0.5 (disjointed proportionality). (C): Gene pairs fulfilling *ϑ*_e_ *<* 0.2 (emergent proportionality). (D): Gene pairs fulfilling *ϑ*_e_ *>* 0.7. Such cut-offs from below induce a cut-off on *ϑ* and an additional restriction on the difference between weighted group variances.

**Figure 3:**
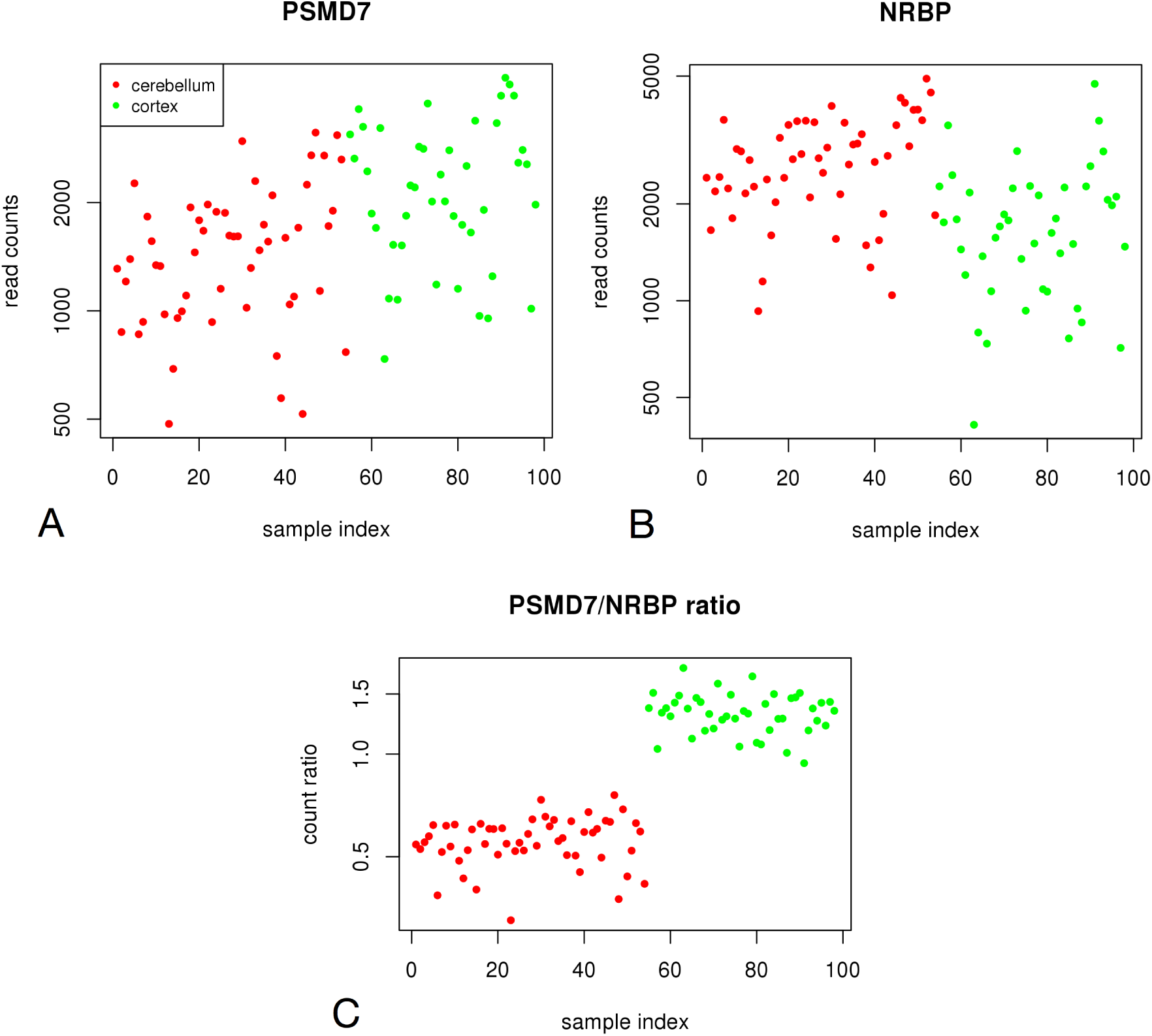
Differential expression of individual genes is not necessary for the pair to be differentially proportional: (A) Read counts plotted against the sample index for the gene PSMD7 (a proteasome subunit). Read counts do not indicate any apparent differences between tissues. (B) A similar situation as in panel A, but for a nuclear receptor binding protein. (C) The ratio plot of the genes from panels A and B. There is a clear difference in the gene ratios, although the individual read counts show no apparent differential expression.

## 2 Methods and Results

### 2.1 Simple statistics for differential proportionality

We start by introducing a short-hand notation which allows us to denote projections of the log-ratios of two vectors **x**, **y** having *n* components (e.g. a gene or transcript pair) onto a subset of size *k*:

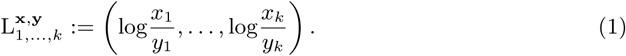

Equivalently, the log-ratio mean (LRM) and variance (LRV) evaluated on this subset are denoted by 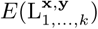 and 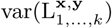 respectively. Let us now assume we have a natural partition of our *n* samples into two subsets (conditions, or tissues) of experimental replicates of sizes *k* and *n-k*. To avoid clutter, we drop **x**, **y** from the notation in the following equation. It is well known that variance evaluates to

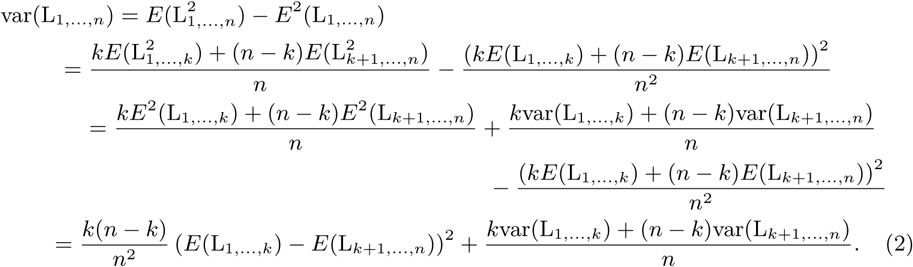

This is the well-known decomposition into between-group variance (first term) and within-group variance (second term) known from analysis of variance (ANOVA). Note that all variances throughout the text are defined as the biased estimators (so the sum of squares are divided by *k* rather than *k -* 1, with *k* the number of summands). As will be seen from the discussion below, differential proportionality can be studied relative to LRV and there is no need for evaluation of the total size of LRV (which is a problem when studying proportionality across all the samples). If we divide (2) by var(L_1,*…,n*_), we obtain as summands the various proportions of (weighted) group variances and of the between-group variance to the overall variance. For illustration, this is visualized as a ternary diagram in Figure 2A. The proportion of within-group variance with respect to overall variance is thus a function of the three LRVs:

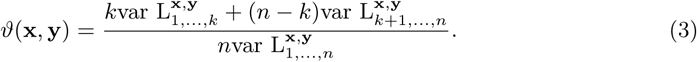

Conveniently, *ϑ* is a number between zero and one. When approaching zero it indicates that the total LRV is explained by the squared difference in group LRMs (Fig. 2B). A large enough difference means that scatter plots of **y** vs. **x** will have different slopes depending on the condition the samples come from. This case is thus characterized by tissue-specific proportionality factors (or group LRMs). We call this type of differential proportionality *disjointed* proportionality here.

We can use *ϑ* for testing this property on our vector pairs and evaluate its significance using a simple permutation test for an estimate of the false discovery rate (FDR). Alternatively, a classical test-statistic known from one-way ANOVA with two groups is the squared *t*-statistic *F*. It is related to *ϑ* by

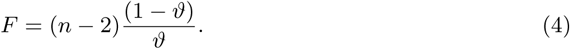

This statistic can be used to do a classical *F*-test of the null hypothesis of equal group (population) LRMs under standard ANOVA assumptions. Note that regardless of the statistic used, multiple testing corrections are especially important in the ratio context due to the large number of gene pairs that get tested. These can be efficiently obtained by estimating the FDR, such as by using the plug-in estimate from a permutation procedure, see e.g. (Hastie *et al.*, 2009).

We have seen that disjointed proportionality describes pairs where between-group variance constitutes the major part of their LRV. Another type of differential proportionality can be defined for those pairs where one of the group LRVs dominates the total LRV. A scatter of **y** vs. **x** will then show proportionality for samples in one condition but no correlation for the other condition. We will call this type of proportionality *emergent* to distinguish it from disjointed proportionality. In complete analogy to the definition of *ϑ*, from (2) we get

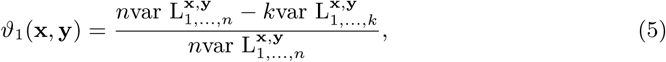

as the proportion of the sum of between-group variance and the LRV of group 2 to the total LRV. Small values of *ϑ*_1_ indicate that the LRV of group 1 constitutes the major part of the total LRV, which is our defining feature of emergent proportionality in group 2. A convenient measure for detecting emergent proportionality regardless of group can be defined as

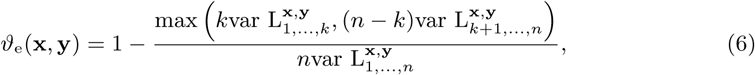

of which a cut-off from above will give us the a set of pairs that are proportional in just one of the two conditions (Fig. 2C). Let us now look at the relationship between *ϑ*_e_ and *ϑ*. Note that we have

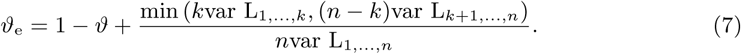

It follows that

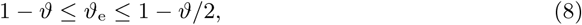

with the equality 1 *- ϑ* = *ϑ*_e_ holding if one of the group LRVs vanishes and *ϑ*_e_ = 1 *- ϑ/*2 in the case of equality of weighted group LRVs *k*var L_1,*…,k*_ = (*n - k*)var L_*k*+1,*…,n*_. It transpires that *ϑ*_e_ can be used to study both types of differential proportionality since large values of it enforce small *ϑ*. For this, a second cut-off on *ϑ*_e_, this time from below, needs to be determined. However, note that a cut-off *ϑ*_e_ *> C* would enforce a somewhat stricter definition on disjointed proportionality, where the induced cut-off *ϑ <* 2(1 *- C*) can only be attained for equality of weighted group LRVs, a condition that is relaxed when going further down with *ϑ*. In fact, cut-offs from below on *ϑ*_e_ cut the upper corner of the ternary diagram with two lines that yield a diamond shape as opposed to the triangle that results from a cut-off on *ϑ* (Fig. 2D). Thus *ϑ*_e_ allows for better control of the correlation within the groups. This can be useful when filtering out those differentially proportional pairs that consist of genes having differential expression but which are not proportional within the groups. This case will be discussed in section 2.4.

### 2.2 Introducing precision weights

RNA-seq data show a pronounced mean-variance relationship that leads to biases when linear models are fit to them. However, log-ratios do not show the mean-variance relationship of the counts directly. The problem here is rather that we should have less confidence in ratios when they involve low counts, as their precision will be lower due to the mean-variance relationship. It has been suggested that an incorporation of the mean-variance relationship via precision weights makes count data accessible for linear modelling (Law *et al.*, 2014) and weighting in general leads to better benchmark performance (Liu *et al.*, 2015). Here we need weights for log-ratios rather than log counts. We can combine the weights *ω*(*x*_*i*_) for read counts of gene **x** in condition *i* into a ratio weight by simply multiplying the weights of both genes involved. Let us denote these weights by

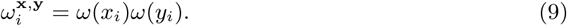

The overall weight of a given ratio for the set of samples 1, *…, k*_1_ from condition 1 is then

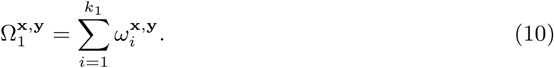

Let us now drop the upper indices for the gene pair. The weighted log-ratio means and variances for a given gene pair in condition 1 will then be

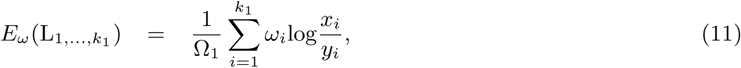

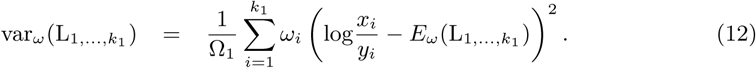

The decomposition of weighted log-ratio variance goes through as before, and a weighted statistic

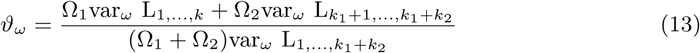

can be defined in analogy to (3). Here we were just interested in the sums, not the actual variances. Note that we can define a unbiased weighted variance estimator specifically for reliability weights. For this, the prefactor in (12) changes from 1*/*Ω_1_ to 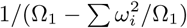.

### 2.3 A moderated statistic for ratios

It has been shown that similarities in expression between the genes can be exploited by assuming an underlying prior distribution of within-group variances and log-fold changes in a gene-expression matrix (Lönnstedt & Speed, 2002; Smyth, 2004). The resulting hierarchical model can be used to derive a moderated *t*-statistic whose parameters can be estimated from the data in empirical-Bayes fashion. The moderated statistic has been shown to be much more powerful than the classical *t*-statistic in simulation-based benchmarks, see (McCarthy & Smyth, 2009). Its effect is especially relevant for small numbers of samples. The moderation shrinks the within-group variance of a gene toward a prior variance and should have a similar effect as the regularization (Witten & Tibshirani, 2009) of a covariance matrix. Here we make use of the gene-wise hierarchical model to moderate the gene *ratio* variances. This approach, although somewhat ad-hoc, is justified by the fact that ratios with unchanged (reference) genes in the denominator are proportional to absolute abundances, which the gene-wise hierarchical model is designed for. The changes of model parameters between arbitrary references are found to be small and are neglected in our approach.

Let us denote the pooled within-group variance of the log-ratios with unchanged reference **z** by

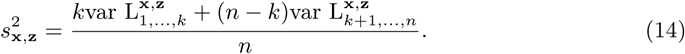

Given the hierarchical model, it was shown that the posterior mean of the inverse population variance *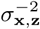*, given the sample variance (14) has the form

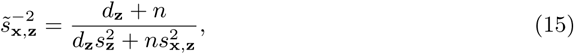

where *d***_z_** and 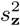
are the parameters of the Gamma distribution serving as a prior for the variance (14). We will not go into more detail of the underlying Bayesian model here but just mention that a moderated *t*-statistic can be obtained by replacing 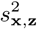 in the original *t*-statistic by 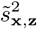. In the following we use (15) as a justification for moderating the within-group variances. This can also be seen as a kind of regularization of the covariance matrix of the log-ratios that have **z** as a reference. From (15) we now derive moderated versions of *F* and *ϑ* for *all* the gene ratios. Let us denote by *F′* the ratio of between-group over within-group LRV for a given gene pair. *F′* is the same as *F* in Equation (4) without the factor (*n -* 2). We have

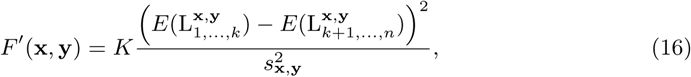

where we also used the short-hand expression

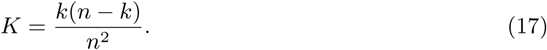

The idea is now to replace the term 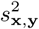 by its moderated version derived from (15). It would only slightly change the parameters (and would lead to loss of symmetry between **x** and **y**) to use a different hierarchical model for each reference *y*. We thus choose a generic reference *z* for obtaining the prior variance. For the moderated *F′* we then find

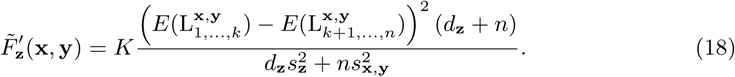

We can further simplify this expression using two relationships following from equations (2) and (3), namely

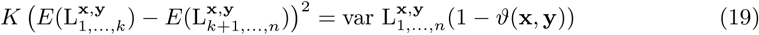

and

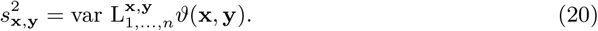

With these, (18) becomes

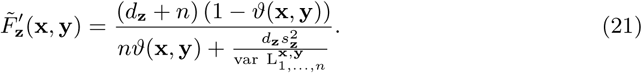

The expression 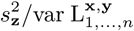 occurring here can be considered the prior *ϑ* to which the original *ϑ*(**x**, **y**) is shrunk. The parameters *d***_z_** and 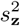 can be determined, e.g. using the limma package (Smyth, 2005). Whether the dependence on the choice of **z** is of any practical importance needs to be investigated empirically. From 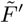 we get immediately the corresponding expressions for 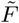 and 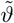 by applying (4):

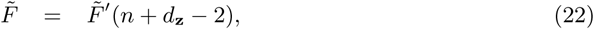

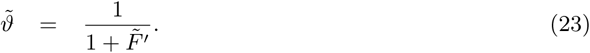

Here, we did not use the weighted variances for clarity and to ease the notational burden; they can be derived in similar fashion.

### 2.4 Relation to differential expression

If we assume that we know the identity of an unchanged reference **z**, it provides us with an ideal normalization (as mentioned in the previous section). The statistic *ϑ*(**x**, **z**) could then be used as a measure for the amount of differential expression of gene **x**, whose log-fold change would be

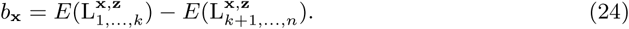

Likewise, within-group variances of individual genes **x**, with an ideal reference **z** can be written as the within-group variances of the ratios with **z**, i.e.

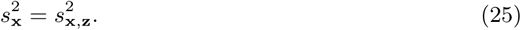

We will now show that if we have two sufficiently strong differentially expressed genes whose log-fold changes have opposite signs, then they will form a differentially proportional pair. Hence, no within-group correlations of the genes are required in this case for their *ϑ* to be small^2^. More formally, we assume

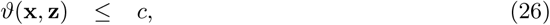

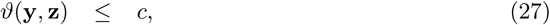

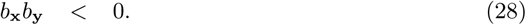

The log-ratio change of the gene pair **x**, **y** is

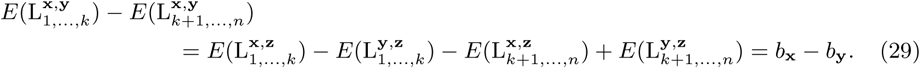

Likewise, the within-group variance of the gene pair **x**, **y** can be written as

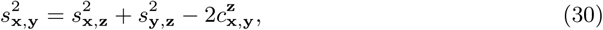

where the term 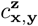 denotes the within-group covariance between **x** and **y** defined by

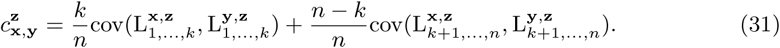

Using (29),(30) and (16) we obtain

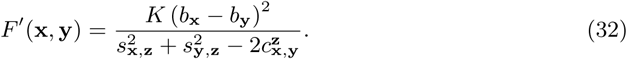

Let us now find a bound from above for the denominator. The definition (16) implies that

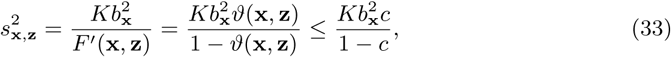

with the bound following from our condition (26), and for 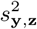 from (27). For an upper bound on the denominator in (32), we can then use

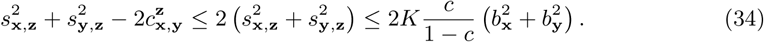

The first bound follows from the fact that the absolute value of the correlation coefficient 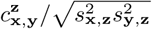 is smaller than one and the arithmetic mean bounds the geometric mean in its denominator. The second bound uses (33). Inserting this back into (32) we find

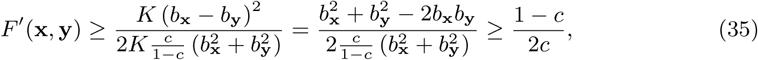

with the last bound following from (28). We thus find that (26)-(28) imply differential proportionality in the sense that

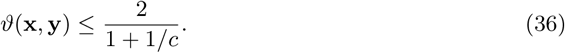

In a similar fashion, more complicated relationships could be derived where the conditions (27) and (28) get relaxed. Instead, we will now look at the reversed question: What can we know about differential expression of the individual genes when the pair is differentially proportional? The only assumption we make is

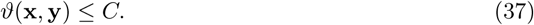

Starting from (32), we have

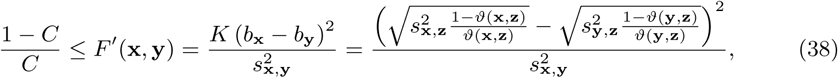

where the last equality was obtained rewriting the second equality in (33). The *ϑ*(**x**, **z**) for which we get the smallest value of *F′* permitted by *C* (i.e. where the equality holds) is obtained by solving the quadratic equation. We get

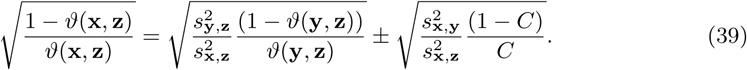

Values of the left-hand side leading to bigger *F′* are obtained below the “*-*” and above the “+” solution. We are in the latter regime if 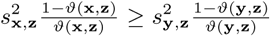. We can assume this to be fulfilled (because **x** and **y** indices can just be swapped in case it is not). Thus choosing the more convenient of the two *ϑ*, we obtain

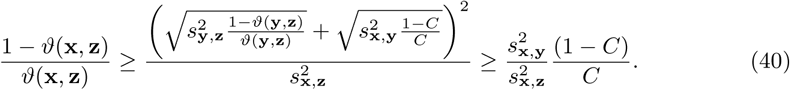

We have thus found the following bound for one of the genes in the gene pair:

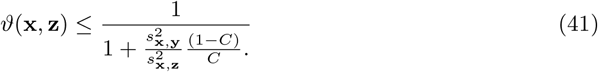

Intuitively this makes sense: when the genes are correlated within the groups, the within-group LRV of the pair 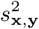 can be small compared to 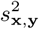, and then *C* may not be sufficiently small for differential expression of **x** (see Figure 3 for an example). For differential expression we thus require a minimum within-group LRV of the differentially proportional pair. Note, however, that although we can control for both 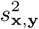 and *C*, the within-group variance of the gene 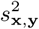 remains inaccessible to us from a strict ratio point of view because it would require our knowledge of the reference **z** leading to the correct normalization. Although for this reason we cannot precisely quantify how small *C* needs to be, the obtained bound on *ϑ*(**x**, **z**) shows qualitatively that differentially proportional pairs with sufficiently high within-group variance will contain at least one differentially expressed gene.

### 2.5 Handling zeros

As reviewed in (Martin-Fernandez *et al.*, 2011), zeros resulting from undersampling (known as count zeros, and a major source of zeros in RNA-seq data) can best be dealt with assuming a Dirichlet prior leading to posterior counts where pseudocounts are added to the original counts. Along the same lines, one can also choose a resampling strategy, where repeated drawings from the posterior distribution lead to a kind of pseudo-replicates that do not contain zeros, which will represent variation expected from the original counts (Fernandes *et al.*, 2013; Tarazona *et al.*, 2015). Since an additive modification does not preserve ratios, a kind of multiplicative modification of a given count

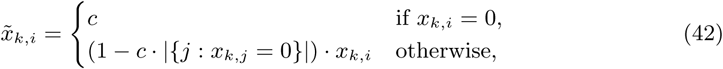

was suggested (Martin-Fernandez *et al.*, 2011). Here the column indices *i* go over the genes in the given condition *k*, and the 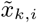 are the counts modified by the pseudocount *c* (which, for simplicity, we assume to be independent of the samples here). The fact that ratios are not preserved when simply adding the pseudocount, however, is felt strongest in the case of low counts, where ratios should not be trusted anyway. To alleviate the problem, it thus seems essential to use the precision weights of section 2.2 when calculating the relevant statistics.

While pseudocounts need an associated distributional theory to estimate them, a well-founded heuristic that has been used widely in data analysis are power transformations of the Box-Cox type. In the limiting case of a power tending to zero, these return the logarithm:

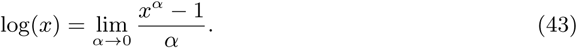

It has been shown by (Greenacre, 2009) that this transformation establishes a connection between Correspondence Analysis (CA) of the transformed data and log-ratio analysis, which is obtained as a limiting case of CA when letting *α* tend toward zero. This is interesting because CA handles zeros naturally. We will briefly describe this replacement strategy here. As shown in (Greenacre, 2011), from re-writing LRV in the form

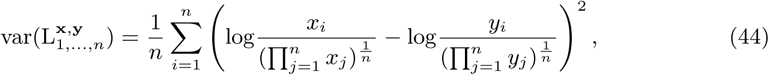

a similarity with the (squared) *χ*^2^ distance used in CA becomes evident. Here we show this distance for data raised to the power of *α* and with rows summing to one:

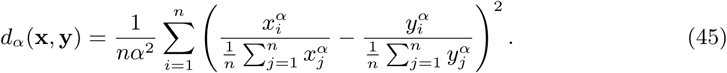

We can obtain (45) directly from (44) by applying (43) for nonzero *α* and replacing geometric by arithmetic means (which is justified in the limit *α →* 0).

A precision-weighted *ϑ* like in (13) that can also handle zeros can thus be defined by

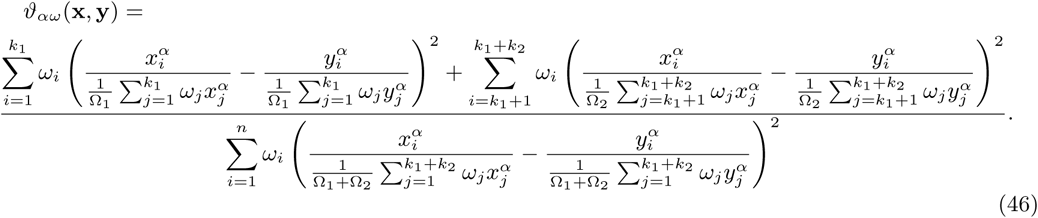

Note that the weighting scheme differs from the one used in CA where weights are determined from row and column sums and low counts get upweighted. The choice of *α* needs to trade off closeness to the original LRV values (for gene pairs not containing zero counts small *α* are more accurate) with the amount by which zeros should get punished (pairs containing zeros can have lower *ϑ* if *α* is larger).

### 2.6 GTEx data

For the practical examples shown here, we used data from the Genotype Tissue Expression (GTEx) project (Lonsdale *et al.*, 2013). Reads were mapped using TopHat2 (Kim *et al.*, 2013) and gene counts were obtained from the Flux Capacitor (Montgomery *et al.*, 2010). 10,842 genes with nonzero counts throughout 7867 samples from 40 tissues were used, then samples were additionally filtered for low ischemic times. Finally, only samples from two approximately balanced brain tissues (54 cerebellum and 44 cortex samples) were retained to match the use case discussed in this article. At an FDR of 5% (estimated by permutation tests) we find a cut-off *ϑ <* 0.94 covering 26.6 million gene pairs (45% of all pairs). At *ϑ <* 0.69 (4.56 million pairs) no false positives were detectable anymore. For high confidence disjointedly proportional pairs with clear within-tissue correlations, we settled for a much stricter cut-off of *ϑ ≤* 0.2 (chosen subjectively by visual inspection of scatter plots) comprising 13,000 pairs. Conventional differential expression analysis using edgeR (Robinson *et al.*, 2010) and DeSeq2 (Love *et al.*, 2014) find about half of all considered genes differentially expressed at an FDR of 5%.

## 3 Outlook

While here we have presented how differential expression of ratios can be formalized, a practical proof of concept needs more in-depth analysis of relevant biological data sets. Preliminary results show that the approach holds great promise since the phenomenon of stoichiometry switches appears to be wide-spread both between tissues and between developmental stages when using data from BrainSpan (Sunkin *et al.*, 2012) (see http://developinghumanbrain.org). These results will be reported elsewhere. The principle is not limited to providing a list of interesting gene pairs. Differential proportionality induces a distance measure between genes (e.g. in the form of *ϑ*) that can be used in a network analysis that is independent of normalization. Our R implementation, available as an addendum to the propr package (Quinn *et al.*, 2017), provides an entry point to relevant graph-based analyses.

## Acknowledgements

I.E. thanks Christian Stenvang for checking differential expression using the edgeR and DeSeq2 packages. T.Q. thanks Tamsyn Crowley and Mark Richardson for their advice and expertise on next generation sequencing. I.E. and C.N. were supported by CRG internal funds provided by the Catalan Government.

## Footnotes

1 This paper is a slightly revised version of the conference paper presented at CoDaWork 2017, the 7th Compositional Data Analysis Workshop, Abbadia San Salvatore, Italy. Its main change is a modified definition of the moderated statistic.

2 This means there are at least two kinds of pairs with small *ϑ*: the ones where genes are proportional within the two groups of samples, and those where both genes are unrelated but differentially expressed individually. The latter have a larger within-group LRV and thus need to compensate with a larger overall LRV.

## References

Bar-Shira, Ossnat, Maor, Ronnie, & Chechik, Gal 2015. Gene expression switching of receptor subunits in human brain development. PLoS computational biology, 11(12), e1004559.

Dillies, Marie-Agnés, Rau, Andrea, Aubert, Julie, Hennequet-Antier, Christelle, Jeanmougin, Marine, Servant, Nicolas, Keime, ćeline, Marot, Guillemette, Castel, David, Estelle, Jordi, et al. 2013 A comprehensive evaluation of normalization methods for Illumina high-throughput RNA sequencing data analysis. Briefings in bioinformatics, 14(6), 671–683.

Erb, Ionas, & Notredame, Cedric 2016. How should we measure proportionality on relative gene expression data? Theory in Biosciences, 135(1-2), 21–36.

Fernandes, Andrew D, Macklaim, Jean M, Linn, Thomas G, Reid, Gregor, & Gloor, Gregory B 2013. ANOVA-like differential expression (ALDEx) analysis for mixed population RNA-Seq. PLoS One, 8(7), e67019.

Greenacre, Michael. Power transformations in correspondence analysis. Computational Statistics & Data Analysis, 53(8), 3107–3116.

Greenacre, Michael 2011. Measuring subcompositional incoherence. Mathematical Geosciences, 43(6), 681–693.

Hastie, Trevor, Tibshirani, Robert, & Friedman, Jerome 2009. The elements of statistical learning: data mining, inference, and prediction. Springer New York.

Kim, Daehwan, Pertea, Geo, Trapnell, Cole, Pimentel, Harold, Kelley, Ryan, & Salzberg, Steven L. 2013 TopHat2: accurate alignment of transcriptomes in the presence of insertions, deletions and gene fusions. Genome biology, 14(4), R36.

Law, Charity W, Chen, Yunshun, Shi, Wei, & Smyth, Gordon K. 2014 Voom: precision weights unlock linear model analysis tools for RNA-seq read counts. Genome biology, 15(2), R29.

Liu, Ruijie, Holik, Aliaksei Z, Su, Shian, Jansz, Natasha, Chen, Kelan, Leong, Huei San, Blewitt, Marnie E, Asselin-Labat, Marie-Liesse, Smyth, Gordon K, & Ritchie, Matthew E. 2015 Why weight? Modelling sample and observational level variability improves power in RNA-seq analyses. Nucleic acids research, 43(15), e97–e97.

Lönnstedt, Ingrid, & Speed, Terry 2002. Replicated microarray data. Statistica sinica, 12, 31–46.

Lonsdale, John, Thomas, Jeffrey, Salvatore, Mike, Phillips, Rebecca, Lo, Edmund, Shad, Saboor, Hasz, Richard, Walters, Gary, Garcia, Fernando, Young, Nancy, et al. 2013 The genotype-tissue expression (GTEx) project. Nature genetics, 45(6), 580–585.

Love, Michael I, Anders, Simon, & Huber, Wolfgang 2014. Moderated estimation of fold change and dispersion for RNA-seq data with DESeqGenome biology, 15(12), 550.

Lovell, David, Pawlowsky-Glahn, Vera, Egozcue, Juan José, Marguerat, Samuel, & Bähler, Jürg 2015. Proportionality: a valid alternative to correlation for relative data. PLoS computational biology, 11(3), e1004075.

Lun, Aaron TL, Marioni, John C, & Bach, Karsten 2016. Pooling across cells to normalize single-cell RNA sequencing data with many zero counts. Genome biology, 17(1), 75.

Martin-Fernandez, Josep Antoni, Palarea-Albaladejo, Javier, & Olea, Ricardo Antonio 2011. Dealing with zeros. Pages 43–58 of: Compositional data analysis: Theory and applications. John Wiley & Sons, Ltd.

McCarthy, Davis J, & Smyth, Gordon K 2009. Testing significance relative to a fold-change threshold is a TREAT. Bioinformatics, 25(6), 765–771.

Montgomery, Stephen B, Sammeth, Micha, Gutierrez-Arcelus, Maria, Lach, Radoslaw P, Ingle, Catherine, Nisbett, James, Guigo, Roderic, & Dermitzakis, Emmanouil T 2010. Transcriptome genetics using second generation sequencing in a Caucasian population. Nature, 464(7289), 773–777.

Quinn, Thomas, Richardson, Mark F, Lovell, David, & Crowley, Tamsyn 2017. propr: An R-package for Identifying Proportionally Abundant Features Using Compositional Data Analysis. Scientific Reports, 7, 16252.

Robinson, Mark D, McCarthy, Davis J, & Smyth, Gordon K 2010. edgeR: a Bioconductor package for differential expression analysis of digital gene expression data. Bioinformatics, 26(1), 139–140.

Smyth, Gordon K 2004. Linear models and empirical bayes methods for assessing differential expression in microarray experiments. Statistical applications in genetics and molecular biology, 3(1), 1–25.

Smyth, Gordon K. 2005 Limma: linear models for microarray data. Pages 397–420 of: Bioinformatics and computational biology solutions using R and Bioconductor. Springer New York.

Sunkin, Susan M, Ng, Lydia, Lau, Chris, Dolbeare, Tim, Gilbert, Terri L, Thompson, Carol L, Hawrylycz, Michael, & Dang, Chinh 2012. Allen Brain Atlas: an integrated spatio-temporal portal for exploring the central nervous system. Nucleic acids research, 41(D1), D996–D1008.

Tarazona, Sonia, Furió-Tarí, Pedro, Turrà, David Pietro, Antonio Di, Nueda, Maŕıa José, Fer-rer, Alberto, & Conesa, Ana 2015. Data quality aware analysis of differential expression in RNA-seq with NOISeq R/Bioc package. Nucleic acids research, 43(21), e140–e140.

Tesson, Bruno M, Breitling, Rainer, & Jansen, Ritsert C 2010. DiffCoEx: a simple and sensitive method to find differentially coexpressed gene modules. BMC bioinformatics, 11(1), 497.

Witten, Daniela M, & Tibshirani, Robert. Covariance-regularized regression and classification for high dimensional problems. Journal of the Royal Statistical Society: Series B (Statistical Methodology), 71(3), 615–636.

